# Impact of Age on Heroin Intravenous Self-Administration in Wistar Rats

**DOI:** 10.64898/2026.05.05.723054

**Authors:** Michael A. Taffe, Sydney L. Mehl, Yanabel Grant, Sophia A. Vandewater

## Abstract

**Background:** Evidence suggests steeper accelerating opioid-related overdose, and non-medical use rates, in middle aged men in recent years compared with younger cohorts. Little is known about whether this is driven by age-related differences in the effects of opioids compared with socio-cultural factors driving non-medical consumption. Rodent models can be useful for dissociating biological from psychosocial factors, however, only minimal evidence exists on the effects of opioids in middle-age rats.

**Objective:** To determine if the anti-nociceptive and rewarding effects of opioids differ between adult and middle-age rats.

**Methods:** Female and male Wistar rats were obtained in early adulthood and examined across 4 to 11 months of age for nociceptive responses to heroin (0-1.56 mg/kg, s.c.) using a warm-water tail withdrawal assay. Subgroups (N=8 per group) were initiated on intravenous self-administration (IVSA) of heroin at either 5 months or 12 months of age.

**Results:** Anti-nociceptive effects of heroin did not differ across age. Female rats that initiated IVSA in early adulthood or middle-age obtained significantly more infusions of heroin than male rats of the same age during acquisition, and in dose-substitution under a FR1 schedule. Male, but not female, rats that initiated IVSA in middle age self-administered less heroin then rats that initiated in early adulthood; this was observed in acquisition and in dose-substitution.

**Discussion:** This study shows that opioid reward is diminished in middle aged male rats. It also found that middle age rats can be used effectively to model opioid-related outcomes, including drug seeking using the IVSA procedure.

## Introduction

Epidemiological evidence suggests more rapidly accelerating opioid-related overdose rates in middle aged individuals in recent years, compared with younger cohorts. Examination of US deaths which involve drugs from 1999-2017 illustrate the rising trend (Hedegaard et al., 2018) and middle age was associated with increased rate of emergency department visits for opioid overdose (Hasegawa et al., 2014). Overall non-medical use of opioids also increases with age, i.e., an odds ratio 1.61 in 35-49 year olds relative to 18-25 year olds (Moss et al., 2026). It is unclear from epidemiological data whether a middle age liability for adverse effects may arise from differences in the reinforcing properties of opioids versus changes in availability, acceptability or other social factors over time. Controlled animal models are often useful for dissociating socio-cultural factors from biological risks and yet little is known about the reinforcing effects of opioids in middle-age in rats.

The vast majority of substance abuse research in pre-clinical rodent models focuses on adolescent and young adult ages, with the latter most often initiated in, e.g. rats, around post-natal day (PND) ∼90-100. For laboratory rat species in which adolescence is generally recognized as the PND 31 to PND 50 interval, and cognitive aging/dementia models typically start around 21 months of age, this leaves the middle to older adult ranges almost entirely unexplored. It is possible to study opioid self-administration in older rats, indeed female rats will self-administer volatilized heroin at PND 410–429, albeit in a small sample of animals previously trained extensively to self-administer nicotine vapor and without comparison to younger animals (Gutierrez and Taffe, 2024). One brief communication also showed that aged (21-24 month old) Wistar rats intravenously self-administer (IVSA) less morphine at low, but not moderate, training doses in 12 h sessions (Bongiovanni et al., 2021). Other work reported that middle aged (the study used 24 month old Brown Norway / Fischer344 F1 hybrid rats that live up to 36 months) rats are more sensitive to the aversive effects of higher dose morphine on brain reward thresholds compared with young adult rats (Jha et al., 2004). This relatively thin evidence suggests more thorough evaluation is needed to determine if development into the middle age range alters opioid reward in rats.

Given the breadth of physiological changes associated with middle age, the distribution of timing of these events across populations and the nearly complete lack of information on differential drug self-administration in middle age of rats, it is necessary to <u>operationalize</u> the middle age equivalent of, e.g., 50 in a human. The female rat is generally recognized to be middle aged somewhere from 8 to 14 months of age (Rubin, 2000) with 65% displaying irregular cycling (Lu et al., 1994) and evidence of neuroendocrine alterations (Downs and Wise, 2009) consistent with a transition to older adulthood as experienced by women. Relatedly, aging research targeting late life cognitive decline generally starts with 21-24 months of age (Gonzalez-Hedstrom et al., 2021; Rojic-Becker et al., 2021). This study therefore uses 12 months of age as the operational definition of middle age.

The goal of the present investigation was to determine if the propensity to self-administer heroin changes in male and female rats across the early adult to middle-aged developmental range. The acquisition of intravenous self-administration (IVSA) of heroin and dose-substitution of heroin, oxycodone and fentanyl were contrasted in 30-week and one-year old rats. Nociception (using a warm water tail-withdrawal assay) was used to monitor any changes in the involuntary behavioral effects of heroin across age or a period of intravenous self-administration.

## Methods

### Subjects

Male and Female Wistar (N=16 per group; CRL) were received at 11 (male) or 12 (female) weeks of age. Rats were randomly assigned to Cohort 1 or Cohort 2 (N=8 per Cohort/sex) to be differentiated by the age of initiation of intravenous self-administration. The vivarium was kept on a 12:12 hour reversed light-dark cycle (lights out at 0800); studies were conducted during vivarium dark. Food and water were provided ad libitum in the home cage. Procedures were conducted in accordance with protocols approved by the IACUC of the University of California, San Diego and were consistent with recommendations in the NIH Guide (Garber et al., 2011).

### Drugs

Heroin (Diamorphine HCl), oxycodone HCl and fentanyl citrate were dissolved in physiological saline for injection. Heroin was provided by NIDA Drug Supply, oxycodone was obtained from Spectrum Chemical MFG Corp (Gardena, CA) and fentanyl was obtained from Cayman Chemical Company (Ann Arbor, MI).

### Nociception and Body Temperature

Tail withdrawal latencies (from 52°C water) and rectal temperature were assessed prior to injection and 30 and 60 minutes post-injection, as previously described (Gutierrez et al., 2022; Nguyen et al., 2018). The impact of doses of heroin (0, 0.32, 0.56, 1.0, 1.56 mg/kg, s.c.) were assessed with order counter-balanced for 0-1.0 mg/kg and the 1.56 mg/kg dose evaluated last for all animals. This was initiated at 22-23 weeks of age and a 3-4 day interval was imposed between evaluations. Tail withdrawal latency following doses of heroin (0, 0.56, 1.0, 1.56 mg/kg, s.c.) was then re-evaluated in all animals starting at ∼40 weeks of age.

### Surgery and Catheter Maintenance

The Cohort 1 rats were surgically prepared with chronic indwelling intravenous catheters at 27-28 weeks of age using gas anesthesia and sterile procedures, as previously described (Aarde et al., 2015; Nguyen et al., 2019; Nguyen et al., 2021). The Cohort 2 rats were similarly prepared with catheters at 50-51 weeks of age. One female rat from each Cohort was euthanized during recovery from the surgical procedure due to illness that could not be resolved with clinical intervention. Catheter patency was assessed with the administration of 6 mg/kg of the ultra-short-acting barbiturate anesthetic, Brevital sodium (1 % methohexital sodium; Eli Lilly, Indianapolis, IN), i.v.. Three non-patent Cohort 1 female rats were re-catheterized in the opposite vein in a second surgical procedure at 32 weeks of age and after recovery were returned to the experimental sequence. The catheters of five Cohort 2 female rats and one male rat were not patent after 14 sessions of acquisition. Three of these female rats were re-catheterized and returned to IVSA acquisition three weeks after they were determined to be non-patent. Re-catheterization was not successful in the other two females and not attempted in the male.

### IVSA Acquisition

Intravenous self-administration (IVSA) was initiated in two-hour sessions with 60 µg/kg/infusion heroin available on a Fixed Ratio 1 schedule of reinforcement with sessions run 5 days per week. The Cohort 1 (adult initiating) animals started at 29 (female; N=7) or 30 (male; N=8) weeks (PND 206, PND 213) of age and the Cohort 2 (middle age initiating) animals started at 53 (female; N=7) or 54 (male; N=8) weeks (PND 368, PND 375) of age. The three re-catheterized female rats were returned to training conditions on the day corresponding to Session 26 of the animals that remained patent.

### Dose Substitution

#### Cohort 1

During Sessions 16-20 rats were tested with different per infusion doses of heroin (0, 15, 30, 60, 120 µg/kg/infusion) available on a FR1 schedule of reinforcement in a counterbalanced order. This was repeated for doses of oxycodone (0, 30, 60, 150, 300 µg/kg/infusion) on Sessions 21-25 and for fentanyl (0, 0.625, 1.25, 2.5, 5.0 µg/kg/infusion) on Sessions 26-30. Three Cohort 1 female rats underwent re-catheterization, due to failing port connections, and were re-started during Session 26 for the remaining animals and continued for up to 11 sessions after the majority of the other animals had concluded the studies.

#### Cohort 2

During Sessions 16-20 rats were tested with different per infusion doses of heroin (0, 15, 30, 60, 120 µg/kg/infusion) available on a FR1 schedule of reinforcement in a counterbalanced order. This was repeated for doses of oxycodone (0, 30, 60, 150, 300 µg/kg/infusion) on Sessions 21-25 and doses of fentanyl (0.625, 1.25, 2.5, 5.0 µg/kg/infusion) on Sessions 26-30. The three re-catheterized female rats started the heroin dose-substitution on the day corresponding to Session 31 of the animals that remained patent.

### Data Analysis

The number of infusions obtained, percent of responses on the drug-associated lever and tail-withdrawal latencies were analyzed by 2-way ANOVA including within-subjects factors for acquisition Session or Dose condition and a between-groups factor for Sex or Age of Initiation of IVSA, as relevant. Mixed effects analysis was used in any cases where data points were missing for a given subject. In all analyses, a criterion of P<0.05 was used to infer a significant difference. Significant main effects were followed with post-hoc analysis using Tukey (multi-level factors), Sidak (two-level factors) or Dunnett (treatments versus control within group) correction. All analysis used Prism for Windows (v. 11.0.0; GraphPad Software, Inc, San Diego CA).

## Results

### Nociception

Withdrawal latencies for the animals that did not experience IVSA in the interval prior to 40 weeks of age show that while older animals are less sensitive to warm water tail immersion (**Figure 1, upper panels**), heroin anti-nociception was unchanged from young adulthood through middle adulthood in rats. The mixed effects analysis confirmed significant effects in the male [Dose: F (3, 21) = 14.32; P<0.0001, Age: F (1, 7) = 7.45; P<0.05, Interaction: n.s.] and female [Dose: F (3, 21) = 8.716;P<0.001, Age: F (1, 7) = 6.38; P<0.05, Interaction: n.s.] groups. Post-hoc exploration of orthogonal comparisons with the Tukey procedure confirmed age-associated differences after saline injection in male rats and after 0.56 mg/kg injection in female rats. There was no significant effect of Age confirmed in the IVSA groups (**Figure 1, lower panels**), but there was a significant effect of Dose in female [F (3, 21) = 16.65; P<0.001] and male [F (3, 21) = 15.91; P<0.0001] groups. (Only four of the female group started on IVSA as early adults were included at the 40 week timepoint because three of the rats were not suspended for the same interval after IVSA due to completing makeup sessions for the dose-response experiments.) The Tukey post-hoc confirmed that all latency after all doses (across age) differed except 0.56 from 0.0 or 1.0 mg/kg in the male non-IVSA group and that latencies after all doses (across age) differed except 0 from 0.56 mg/kg and 1 from 1.56 mg/kg in the female non-IVSA group.

**Figure 1:**
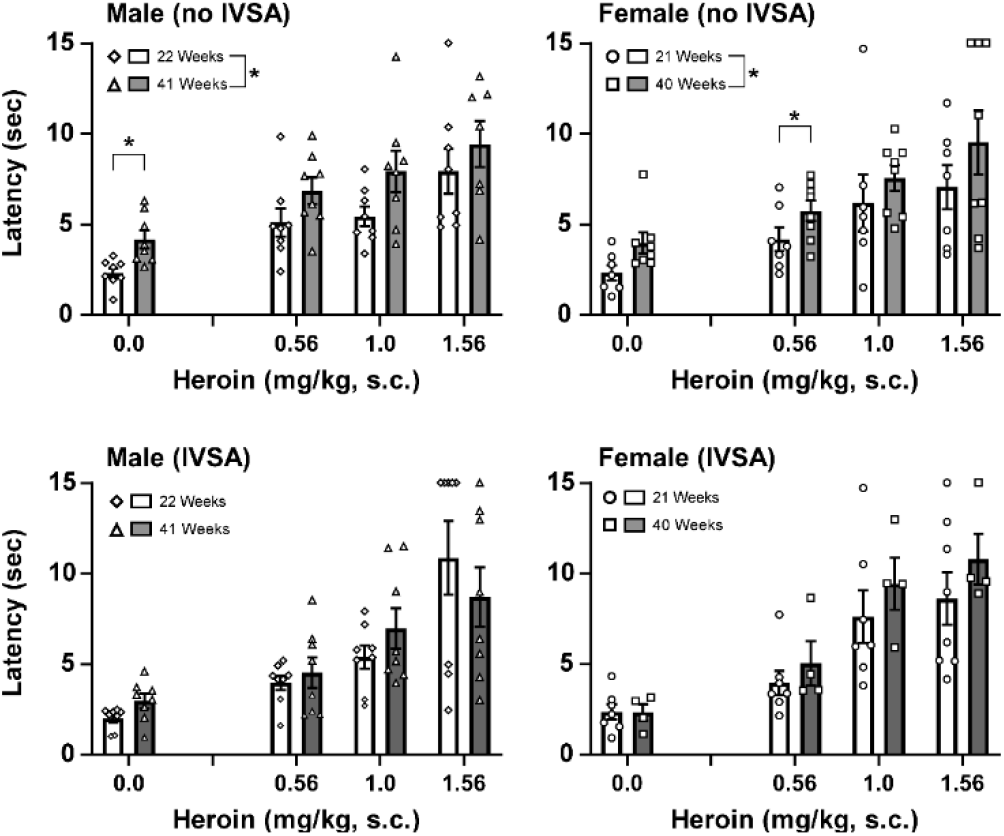
**Upper)** Mean (N=8 per group; ±SEM) and individual tail-withdrawal latency of Wistar rats without IVSA experience in early adulthood and approaching middle age. **Lower)** Mean (N=4-8 per group; ±SEM) and individual tail-withdrawal latency of Wistar rats with IVSA experience between early adulthood and approaching middle age. A significant slowing associated with age is indicated with *.

### Early Adulthood: IVSA Acquisition

Female rats that initiated heroin IVSA in early adulthood self-administered more infusions during the first 15 sessions (**Figure 2**), as confirmed by significant effects of Sex [F (1, 13) = 43.02; P<0.0001], Session [F (4.495, 55.54) = 21.39; P<0.0001] and the Interaction of Sex with Session [F (4.495, 55.54) = 3.31; P<0.05)] The Tukey post-hoc test further confirmed a significant difference between the sexes for all sessions except the 1^st^ and 10^th^. The post-hoc test also confirmed significantly more infusions compared with the first session for female (Sessions 9, 11-15) and male (Sessions 9, 12) groups. There were no sex differences in the percentage of responses on the drug-associated lever confirmed (Sex: n.s., Session: F (6.166, 71.79) = 6.22; P<0.0001, Interaction: n.s.). The Tukey post-hoc test confirmed significantly higher lever discrimination, compared with the first session, for Sessions 8-15 across groups. Three female rats had a port failure (after the 4^th^, 9^th^, and 11^th^ sessions). Four baseline sessions were conducted after re-catheterization and these were included in the analysis as the 12-15^th^ sessions with the other missing sessions omitted.

**Figure 2:**
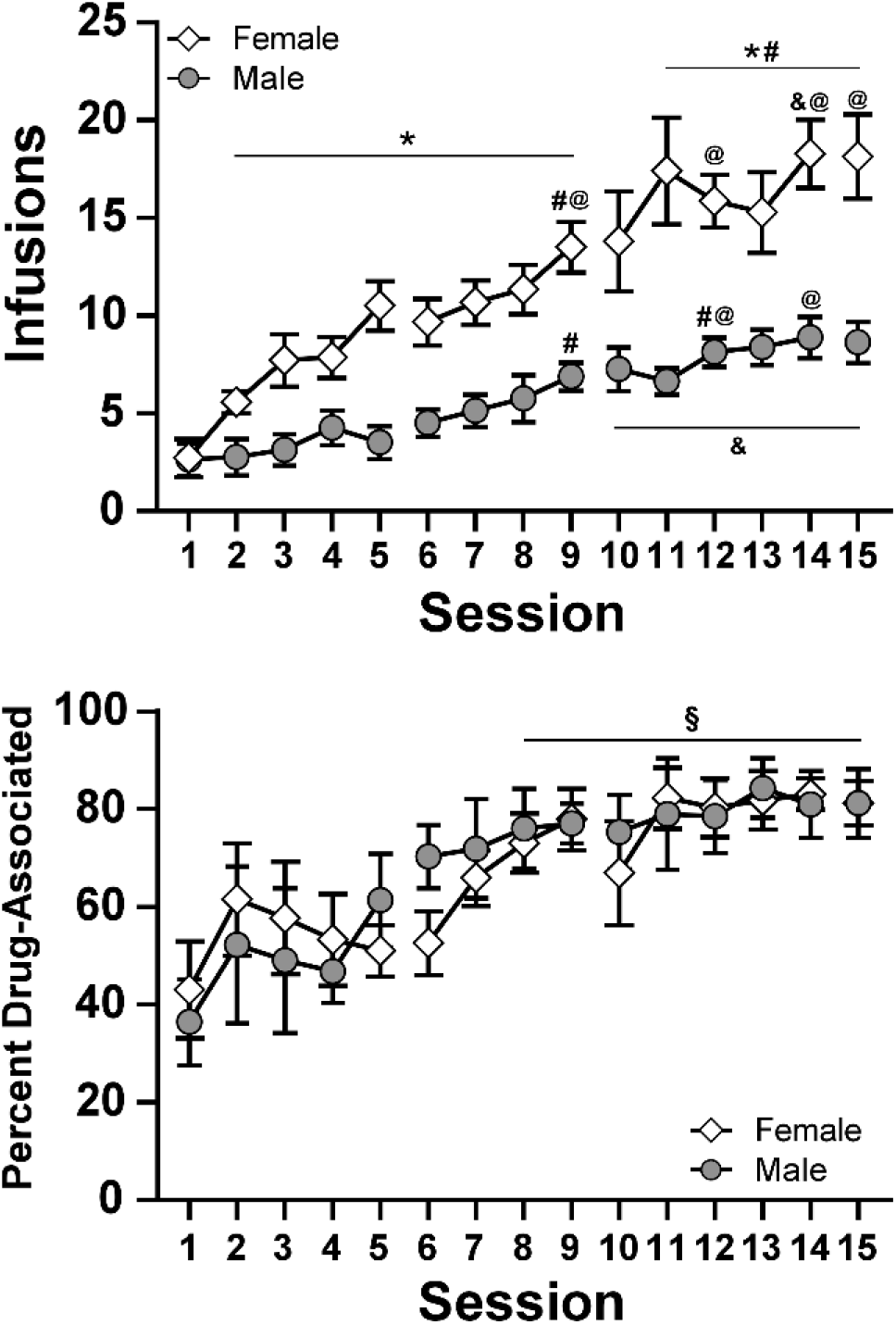
Mean (±SEM) infusions of heroin (60 µg/kg/infusion) obtained by adult (29-30 weeks of age at initiation) male (N=8) and female (N=7) Wistar rats during acquisition, and the percentage of responses on the drug-associated lever. A significant difference between groups is indicated with *, a difference within group from Session 1 with #, from Session 2 with @ and from Session 3 with &. A difference from Session 1, across groups, is indicated with §.

### Middle Age: IVSA Acquisition

Middle age <u>male rats</u> that initiated IVSA at 54 weeks of age obtained fewer infusions of heroin (60 µg/kg/infusion) during acquisition (**Figure 3**) compared with their counterparts who initiated IVSA in earlier adulthood (30 weeks), and the ANOVA confirmed significant effects of Age of initiation [F (1, 13) = 7.95; P<0.05), Session [F (4.463, 58.03) = 9.24; P<0.0001] and the Interaction [F (4.463, 58.03) = 2.58; P<0.05] on infusions obtained. The Tukey post-hoc test limited to orthogonal comparisons confirmed significant group differences on Sessions 6-7, 9-12 and 14-16. Within group, the middle aged animals did not obtain significantly different infusions across any pair of sessions. In contrast, the younger adults obtained significantly more infusions compared with the first session on Sessions 9 and 12, more than Session 2 on Sessions 12 and 14 and more than Session 3 on Sessions 10-15. There were no significant effects of Age of initiation, Session or the interaction on the percentage of responses on the drug-associated lever.

**Figure 3:**
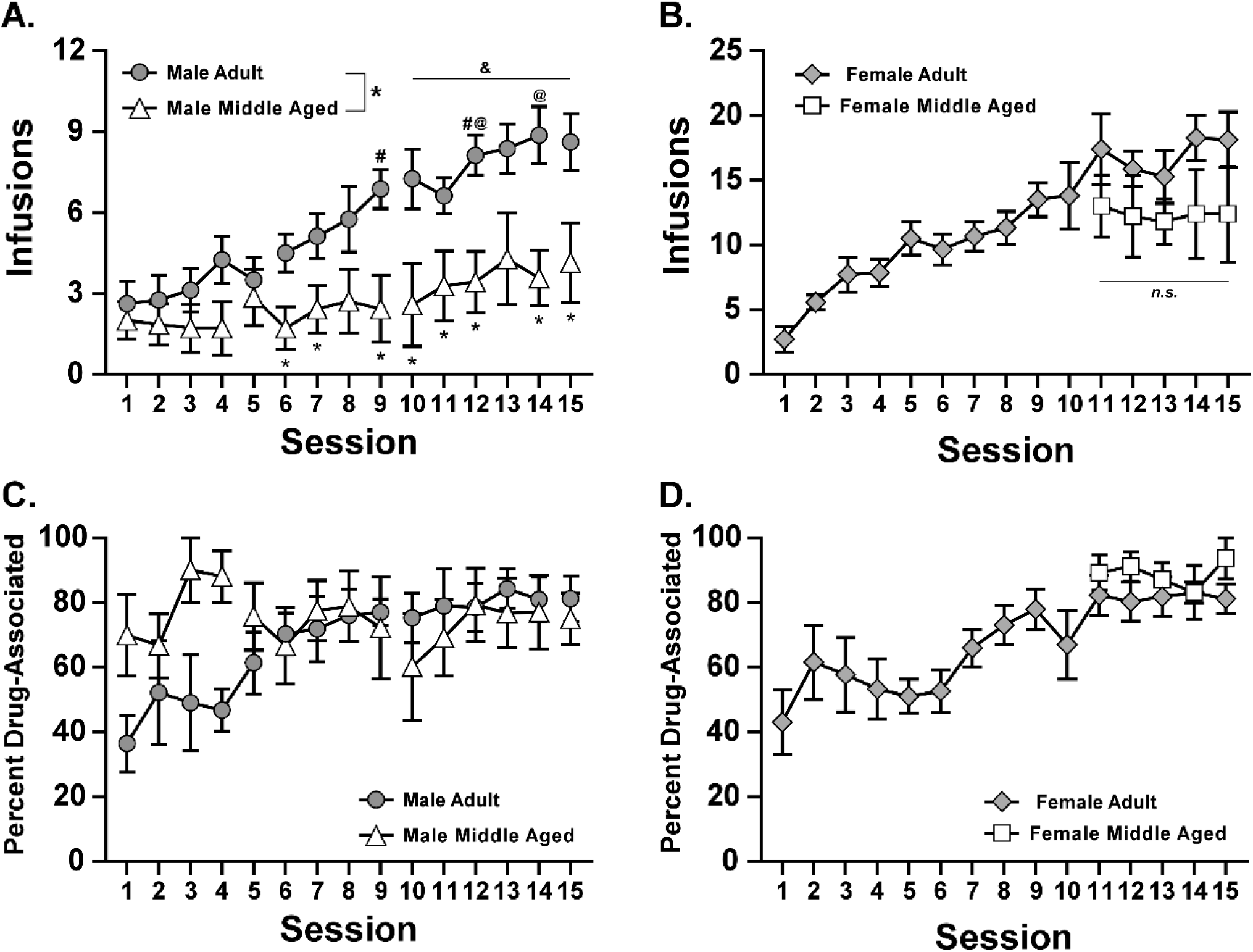
Mean (±SEM) infusions of heroin (60 µg/kg/infusion) obtained by adult (29-30 weeks of age at initiation) and middle aged (53-54 weeks of age at initiation) A) male (N=7-8) and B) female (N=5-7) Wistar rats during acquisition, and C,D) the percentage of responses on the drug-associated lever. N.b. the scale for male and female infusions is different to enhance visual clarity of age-associated differences. A significant difference between groups is indicated with *, a difference within group from Session 1 with #, from Session 2 with @ and from Session 3 with &.

Middle age <u>female rats</u> could not be definitively compared with the younger animals across initial acquisition due to the catheter patency failures in early acquisition. The analysis therefore aligned the final 5 sessions of the two animals with patent original catheters with the five sessions post-recatheterization for three additional animals.

No significant differences in infusions obtained, or in the percentage of responses on the drug-associated lever, were confirmed for the 1 year old and the adult-initiating female groups. Comparison of the final five acquisitions sessions across sex for the 1-year initiating groups confirmed a significant effect of Sex [F (1, 11) = 11.56; P<0.01] on infusions acquired but not on the percentage of responses on the drug-associated lever.

### Heroin Dose Substitution

Female rats in early adulthood self-administered more infusions than males during the heroin dose substitutions under a FR1 schedule of reinforcement (**Figure 4 A**). The mixed effects model confirmed significant effects for heroin infusions [Sex: F (1, 13) = 16.43; P<0.005, Dose: F (2.276, 28.45) = 25.00; P<0.0001, Interaction: n.s.], and the Tukey post-hoc tests limited to orthogonal comparisons confirmed significant sex differences for heroin (0, 15, 60, 120 µg/kg/infusion). The Tukey post-hoc test of the marginal means for Dose confirmed significant differences from Saline (60-120 µg/kg/infusion) and from either 15 or 120 µg/kg/infusion (30-60 µg/kg/infusion). This was also the case for the middle-aged rat groups and the mixed effects model confirmed significant effects on the infusions obtained [Sex: F (1, 11) = 25.33; P<0.0005, Dose: F (2.02, 21.69) = 16.28; P<0.0001, Interaction: F (2.018, 21.69) = 6.17; P<0.01]. The Tukey post-hoc tests limited to orthogonal comparisons confirmed significant sex differences for all heroin doses.

**Figure 4:**
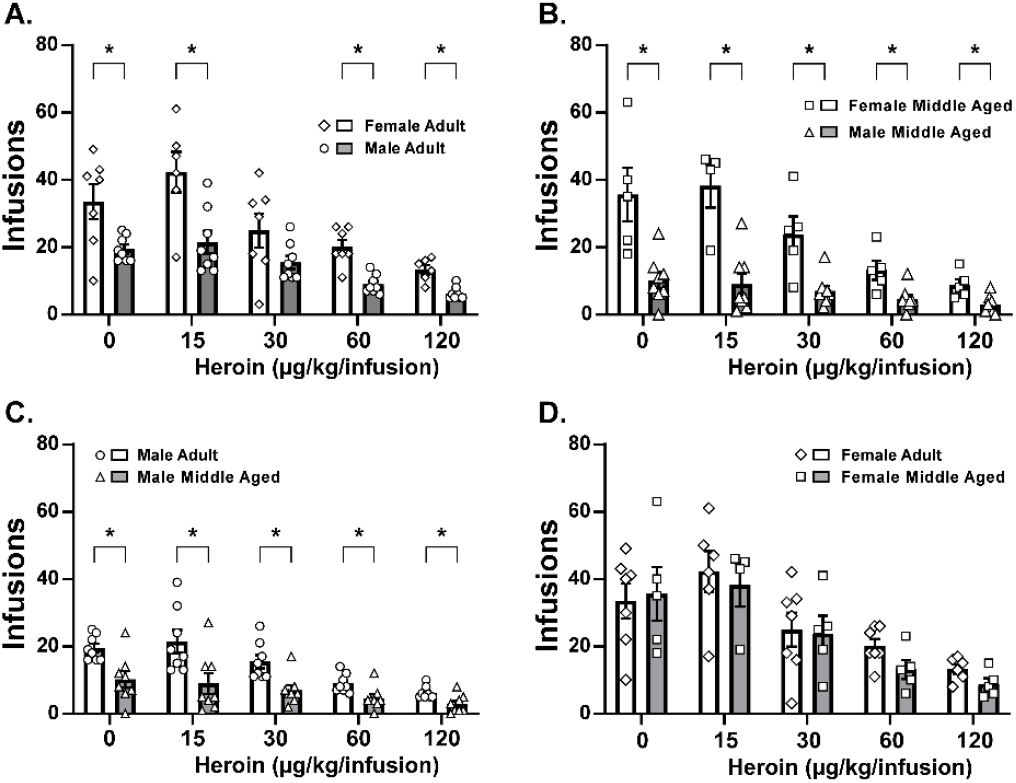
Mean (±SEM) and individual infusions of heroin (0-120 µg/kg/infusion) obtained in a dose-substitution procedure under a FR1 contingency by **A)** female (N=7) and male (N=8) rats which initiated IVSA at 29-30 weeks of age, **B)** female (N=5) and male (N=8) rats which initiated IVSA at 53-54 weeks of age, the male and female **C)** adult and **D)** middle aged initiating groups. A significant difference between groups is indicated with *.

In the age comparison, the middle aged <u>male rats</u> acquired fewer infusions of heroin than did the younger initiating male cohort across a range of heroin doses (**Figure 4 C**) in a post-acquisition dose-response determination under a FR1 schedule of reinforcement as confirmed by a significant effect of Age of initiation [F (1, 14) = 10.95; P=0.01), Dose [F (1.870, 26.17) = 28.79; P<0.0001] and the Interaction [F (1.870, 26.17) = 4.41; P<0.05]). The Tukey post-hoc test limited to orthogonal comparisons confirmed significant group differences on each Dose condition. The middle aged animals obtained significantly fewer infusions of the 120 µg/kg/infusion dose compared with vehicle or 30 µg/kg/infusion. For the early adult initiating cohort, all dose conditions differed from each other condition save that 15 and 30 µg/kg/infusion did not differ from the vehicle condition. The middle aged <u>female rats</u> acquired similar numbers of infusions of heroin across the range of doses (significant effect of Dose F (2.257, 20.88) = 16.20; P<0.001) compared with the younger initiating female cohort (**Figure 4 D**). The Tukey post-hoc test of the marginal means for Dose confirmed significant differences between all dose conditions save that 30 µg/kg/infusion did not differ from saline or 60 µg/kg/infusion and 15 µg/kg/infusion did not differ from saline. See **Supplemental Materials Figures S3, S4** for the oxycodone and fentanyl substitution experiments.

## Discussion

The study shows, first, that male rats that initiate intravenous self-administration (IVSA) of heroin in middle age (>12 months) self-administer fewer infusions of heroin during acquisition compared with male rats that initiate heroin IVSA in early adulthood; however, this pattern was not present for female rats. The second major observation of the study was that female rats obtained significantly more infusions of heroin than did male rats in early adulthood and during middle age. Finally, there was not any major change in the anti-nociceptive effects of heroin across the ∼21-22 week to ∼40-41 week age of Wistar rats, despite a decreased baseline sensitivity to hot water tail immersion in middle aged rats of each sex. Overall, this suggests that while the impact of heroin on nociception is unchanged in each sex, the efficacy of heroin as a reinforcer declines in middle age in male rats.

The early adult initiating female rats obtained significantly more infusions of heroin than their male counterparts during acquisition, with differences observed as early as the second day of drug access (**Figure 2**). This difference was continued in the Fixed-Ratio dose-substitution experiments for heroin (**Figure 4**) as well as oxycodone, but not for fentanyl (**Supplemental Figure S3, S4**). There were no sex differences observed in the proportion of responses directed at the drug-associated lever during acquisition, and both sexes reached group means of >80% in the final three sessions, thereby confirming this was not a learning effect. Although the acquisition protocol was interrupted for the middle-aged females, there was a similar sex difference when stable levels of responding were reached and this difference continued into the heroin (**Figure 4B**) and oxycodone (**Supplemental Figure S3 B**) dose-substitution experiments. Thus, the outcome at both ages is consistent with prior studies where female rats obtained more opioid infusions (Cicero et al., 2003; Deckers et al., 2025; Klein et al., 1997; Nguyen et al., 2020; Taffe et al., 2026; Towers et al., 2022; Townsend et al., 2019), and is inconsistent with opioid IVSA studies in which no sex differences in opioid intake were observed (Carter et al., 2021; D’Ottavio et al., 2023; Lynch and Carroll, 1999; Mavrikaki et al., 2021; Mavrikaki et al., 2017; Rakowski et al., 2025).

Because this study showed that male, but not female, rats who initiate heroin IVSA in middle age self-administer fewer infusions of heroin during acquisition and at every dose in a FR1 dose-substitution procedure (**Figure 4C**), compared with rats that initiated heroin IVSA in early adulthood (∼29-30 weeks of age), the data suggest an interaction between sex and age in terms of the reinforcing properties of heroin. A similar age-associated reduction was observed for self-administration of different doses of oxycodone in the male rats and also the female rats (**Supplemental Figure S3**). As there were no significant differences in the percentage of responses directed at the drug-associated lever during acquisition, differences in learning to associate behavior with the rewarding effects of intravenous heroin are unlikely to explain the age difference. These results are congruent with a prior finding that aged (21-24 month old) rats IVSA less morphine than earlier adult animals (Bongiovanni et al., 2021) in 12 h sessions. These results are discordant with a study showing aged mice (101 weeks) self-administered more remifentanil intravenously compared with young mice (Sharpe et al., 2025), although obviously species differences, and the impact of a very short acting opioid such as remifentanil would require additional work to fully explicate. Other work reported the aversive effects of high dose morphine on brain reward thresholds occurs at lower doses in middle aged (the study used 24 month old Brown Norway / Fischer344 F1 hybrid rats that live up to 36 months) versus young adult rats (Jha et al., 2004). Although it is possible that aversion contributed to age differences in the male rats in this study, the dose-substitutions did not confirm any interaction of Age of initiation with Dose which is inconsistent with such a conclusion.

In addition to the lack of difference in heroin anti-nociception associated with middle age, there was no lasting alteration in the anti-nociceptive effects of heroin caused by the 10-week IVSA protocol, despite what appeared to be an initial increase in sensitivity to the noxious stimulus after vehicle injection, observed several weeks after IVSA. In the female animals, follow-up investigation found *decreased* sensitivity to lower temperature water immersion in the IVSA-experienced group. The lack of a similar IVSA-related difference in baseline nociception of the male rats may be a reflection of their overall lower level of voluntary drug exposure. The heroin challenges suggest, however, that age-related differences in voluntary consumption in the middle-age initiation group were not associated with any age-associated reduction in analgesic efficacy.

There are a few minor limitations to the study. The age of the earlier adult animals was slightly older than is typical for many opioid self-administration studies where initiation from PND 84-120 is common (Ahmed et al., 2000; Carroll et al., 2001; Matzeu and Martin-Fardon, 2020; Mavrikaki et al., 2017; Venniro et al., 2017; Wade et al., 2015); in many cases age has to be inferred from the body weight. Thus, it cannot be ruled out that animals initiated on IVSA at, e.g., PND 100 would differ from the present groups. Nevertheless, it is relatively common for studies that are initiated circa PND 100 to continue with longitudinal manipulations that stretch into the age interval used here for the younger-initiation cohort. In some cases IVSA is initiated weeks into adulthood due to, e.g., a longitudinal pre-IVSA treatment / evaluation design (Nguyen et al., 2021; Nguyen et al., 2018; Pravetoni et al., 2014; Smith and Pitts, 2012). It was not ideal that so many of the female animals lost their catheter patency during acquisition and had to be re-started. This led to an uncertainty about how much heroin they were exposed to via intravenous delivery versus subcutaneous leak of the catheter and about the consequences of the 3-week interval from discontinuation to re-start of IVSA.

In conclusion, the study found that opioid reward is diminished in middle aged male rats, but this did not extend to female middle aged rats. It also confirmed that female rats self-administer more infusions of heroin and other opioids on a weight-adjusted basis from early adulthood to middle age. In the larger sense, this investigation shows that middle age rats can be used effectively to model opioid-related outcomes, including drug seeking using the IVSA procedure.

## Acknowledgements

The authors are grateful to Helen Kim, Christianne J. Perral and Sara Rahman for technical assistance.

