## Supplemental Materials for "Impact of Age on Heroin Intravenous Self-Administration in Wistar Rats"

**Supplemental Introduction**

Recent efforts to include both sexes of laboratory rats in self-administration studies have produced a diversity of outcomes with respect to sex-differences in opioid IVSA. In many cases no general differences are found (Barattini et al., 2024; Carter et al., 2021; Lynch and Carroll, 1999; Maitland et al., 2024; Mavrikaki et al., 2021; Mavrikaki et al., 2017; Nguyen et al., 2024; Rakowski et al., 2025), in some cases female rats obtain more drug infusions (Cicero et al., 2003; Deckers et al., 2025; Nguyen et al., 2020; Townsend et al., 2019) and in limited cases, males obtain more drug (Rakowski et al., 2025). No sex differences for morphine IVSA were reported in aged rats (Bongiovanni et al., 2021). Given the diversity of rat strains, self-administration models and the opioid identity it can be difficult to ascertain the conditions which do, and do not, reliably show sex differences. Thus, it remains critical to include both sexes where possible.

**Supplemental Methods**

Surgery and Catheter Maintenance:

As previously described (Aarde et al., 2015; Nguyen et al., 2019; Nguyen et al., 2021), catheters consisted of a 14-cm length polyurethane based tubing (MicroRenathane®, Braintree Scientific, Inc, Braintree MA, USA) fitted to a guide cannula (Plastics one, Roanoke, VA) curved at an angle and encased in dental cement anchored to an ~3-cm circle of durable mesh. The catheter tubing was passed subcutaneously from a port at the back, inserted into the jugular vein and secured gently with suture thread. A liquid tissue adhesive was used to close the incisions (3M™ Vetbond™ Tissue Adhesive; 1469S B). A minimum of 7 days was allowed for surgical recovery prior to starting the experiment. Catheters were flushed with ~0.2-0.3 ml heparinized (32.3 USP/ml) saline before sessions and ~0.2-0.3 ml heparinized saline containing cefazolin (71 mg/mL) after sessions. Catheter patency was assessed with the administration of 6 mg/kg of the ultra-short-acting barbiturate anesthetic, Brevital sodium (1 % methohexital sodium; Eli Lilly, Indianapolis, IN), i.v.. Animals with patent catheters exhibit pronounced loss of muscle tone within ~3 s of infusion. Animals that failed to display these signs were discontinued from the study and any data that were collected after the previous passing of the test were excluded from analysis.

Progressive Ratio Dose Substitution:

In this paradigm, rats are run in sessions with drug infusions available on a Progressive Ratio (PR) schedule of reinforcement whereby the required response ratio was increased after each reinforcer delivery, within a session as determined by the following equation (rounded to the nearest integer): Response Ratio=5e^(injection number*j)–5 (Richardson and Roberts, 1996). In this study the j value was set to 0.2 and sessions were a maximum of 3 h in duration. Cohort 1 rats were switched to 3-h sessions with drug infusions available on the Progressive Ratio schedule after completion of the fentanyl dose-response under the FR1 contingency. Doses of heroin (15, 30, 60, 120 µg/kg/infusion; Sessions 31-34), oxycodone (30, 60, 150, 300 µg/kg/infusion; Sessions 35-38) and fentanyl (0.625, 1.25, 2.5, 5.0 µg/kg/infusion; Sessions 39-42) were assessed in a counterbalanced order within drug. The PR was not tested in the middle aged rats due to observing no difference in breakpoints associated with dose in the early adult animals and therefore not having a clear comparison.

### Supplemental Results

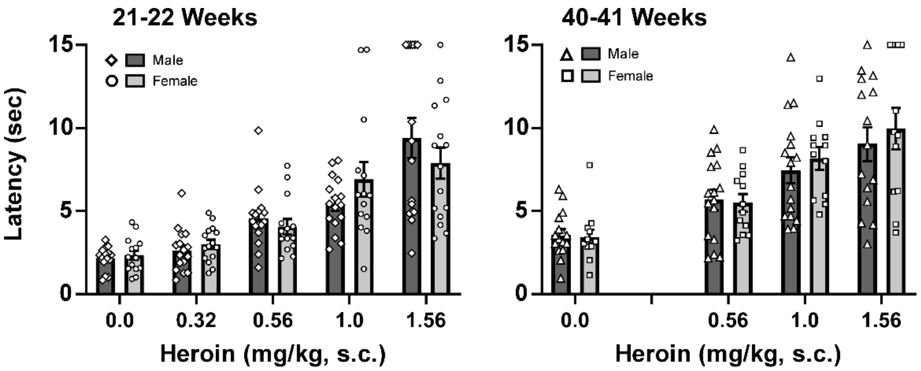


***Figure S1:*** *Mean (N=12-16 per group; ±SEM) and individual tail-withdrawal latency of Wistar rats in early adulthood (N=16 per sex) and approaching middle age (N=12 female, N=16 male).*

<u>Nociception:</u>

Tail withdrawal latencies were significantly slowed by heroin injection in a dose-dependent manner in male and female rats observed at 21-22 weeks of age and at 40-41 weeks of age (i.e., starting 11 days after the last fentanyl PR IVSA session for cohort 1) There were no significant effects of Sex in the full sample at either age (**Figure S1**). The mixed-effects analysis of tail withdrawal latency at 21/22 weeks of age confirmed a significant effect of Dose [F (2.173, 61.38) = 34.58; P<0.0001], but not of Sex or the interaction. The Tukey post-hoc further confirmed that across sex, latency after every dose differed significantly from every other dose condition, save that vehicle did not differ from 0.32 mg/kg and the 1.0 and 1.56 mg/kg dose conditions did not differ from each other. The mixed-effects analysis of tail withdrawal latency at 40/41 weeks of age also confirmed a significant effect of Dose [F (2.252, 57.79) = 29.53; P<0.0001], but not of Sex or the interaction. The Tukey post-hoc further confirmed that across sex, every dose differed significantly from every other, save that the 1.0 and 1.56 mg/kg dose conditions did not differ from each other.

A moderate hyperalgesia was observed in the IVSA cohorts following saline challenge insofar as the sedentary cohort of both male and female rats exhibited slower withdrawal latencies 30 minutes after saline injection compared with the PND143-150 observation, whereas the IVSA groups did not (**Figure S2 A**). However, the Cohort 1 (early adult IVSA) and Cohort 2 (middle age IVSA) groups did not differ significantly from each other at either timepoint. To further examine potential lasting differences, the rats were re-assessed at multiple water bath temperatures under the baseline conditions, i.e. two water bath temperatures were evaluated each day with 60 minutes between evaluations (**Figure S2 B,C**). Tail withdrawal from 46°C and 50°C water was counterbalanced on one day and from 48° and 52°C water on another day. Two female IVSA rats had completed dose-substitutions in IVSA and were included here, thus N=6 for that group.

Analysis confirmed withdrawal latency differed significantly in male [Group: n.s.; Bath Temperature: F (2.372, 33.20) = 8.83; P<0.001; Interaction of Group with Bath Temperature: n.s.] and female rats [Group: F (1, 12) = 10.06; P<0.01; Bath Temperature: F (1.675, 20.10) = 17.90; P<0.0001; Interaction of Group with Bath Temperature: F (1.675, 20.10) = 6.98; P<0.01]. The post-hoc test confirmed a significant difference between female groups at 46°C and 48°C. Tail withdrawal latency differed significantly by bath temperature (**Figure S2 B,C**) in the male (46°C v 52°C, 48°C v 50-52°C) and female (46°C v 48-52°C) groups. There was an inadvertent scheduling error which conflated the IVSA status with the testing order. Thus, the two cooler temperatures were re-evaluated on another test day, with ascending order used for all animals. This confirmed slower latencies in the IVSA group and a significant interaction of bath temp with group in the females was significant at 48°C by the post-hoc test (**Figure S2 D**).

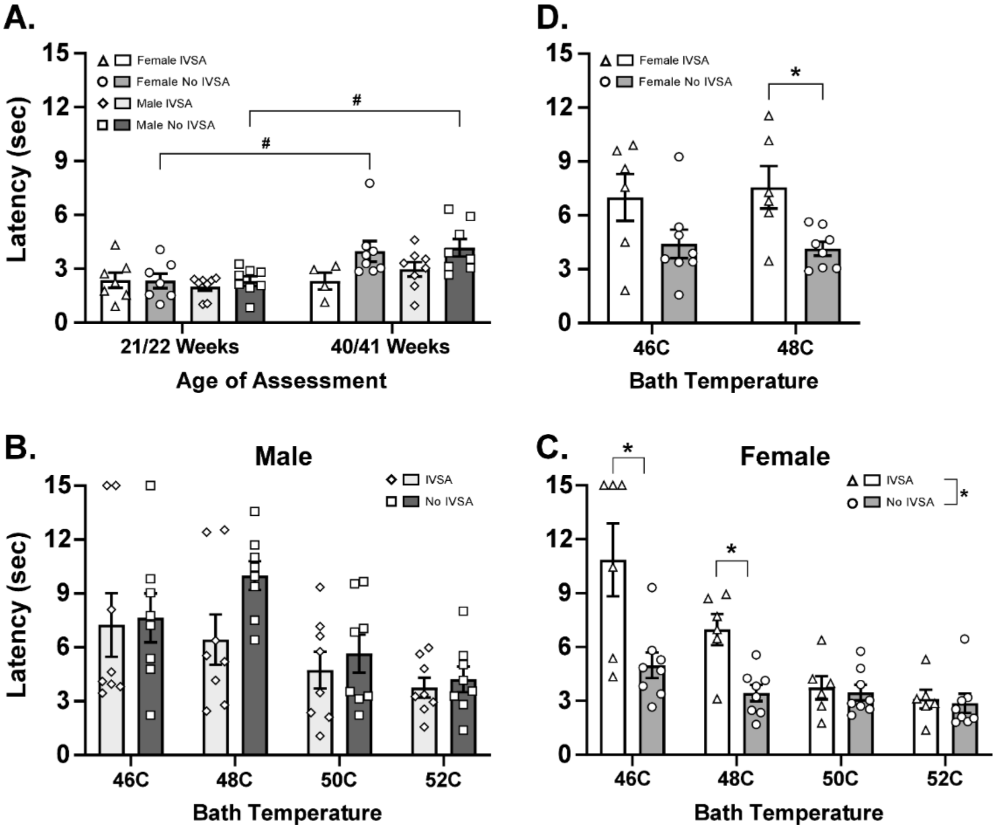


***Figure S2:*** *Mean (±SEM) and individual latency of tail withdrawal from 52°C water observed A) 30 minutes after saline injection from the 21/22 Weeks and 40/41 Weeks dose-effect determinations. Tail withdrawal latencies for B) male and C) female animals in middle age after IVSA training or no training, assessed at four different bath temperatures; N=8 per group except female IVSA N=4 due to two re-catheterized animals catching up on dose-substitutions, two euthanized due to illness prior to completing this study. D) A follow-up assessment of the female animals limited to the two cooler temperatures. A significant difference between groups is indicated with *.*

### Oxycodone Dose Substitution

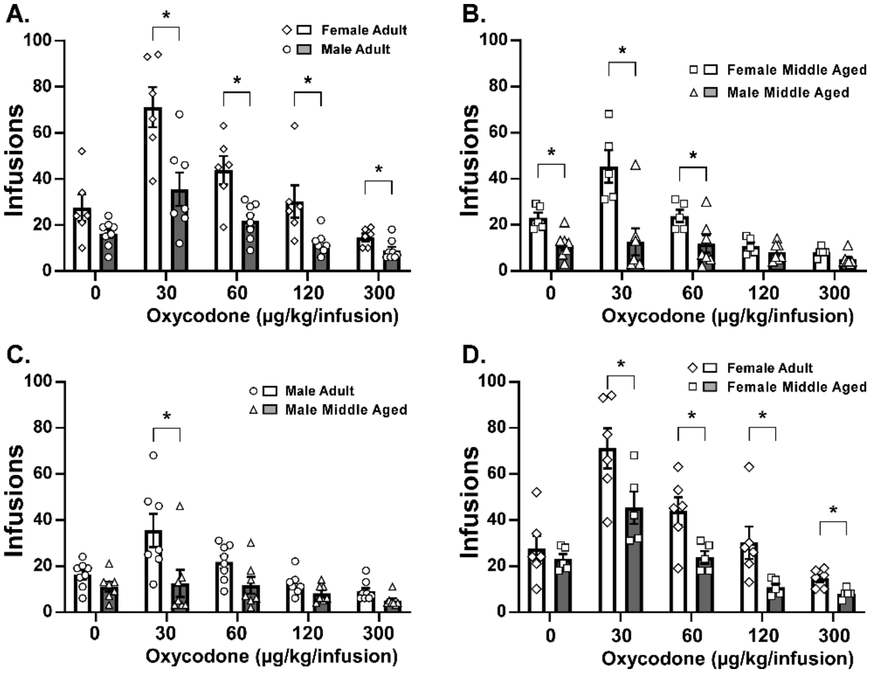


***Figure S3:*** *Mean (±SEM) and individual infusions of oxycodone (0-300 μg/kg/infusion) obtained in a dose-substitution procedure under a FR1 contingency by* ***A)*** *female and male rats which initiated IVSA at 29-30 weeks of age,* ***B)*** *female and male rats which initiated IVSA at 53-54 weeks of age, the male and female* ***C)*** *adult and* ***D)*** *middle aged initiating groups. A significant difference between sexes is indicated with *.*

<u>*Sex differences in early adulthood:*</u>

Female rats in early adulthood (N=6) self-administered more infusions of oxycodone (**Figure S3 A**) than males (N=8) during the dose substitutions under a FR1 schedule of reinforcement. The mixed effects model confirmed significant effects for oxycodone [Sex: F (1, 12) = 33.82; P<0.0001, Dose: F (2.038, 23.94) = 22.33; P<0.0001, Interaction: n.s.] and fentanyl [Sex: F (1.175, 12.34) = 11.93; P<0.005, Dose: n.s., Interaction: n.s.]. The Tukey post-hoc tests limited to orthogonal comparisons confirmed significant sex differences for specific doses of oxycodone (30-300 μg/kg/infusion) and fentanyl (2.5-5.0 μg/kg/infusion). The Tukey post-hoc test of the marginal means for Dose for oxycodone confirmed significant differences from 30 μg/kg/infusion for all other dose conditions and between 60 and 300 μg/kg/infusion. The post-hoc analysis for fentanyl confirmed significant differences from Saline (0.625-1.25 μg/kg/infusion), from 5.0 μg/kg/infusion (0.625-2.5 μg/kg/infusion), and between 1.25 and 2.5 μg/kg/infusion.

<u>*Sex differences in middle age:*</u>

Middle aged female rats (N=5) self-administered more infusions of oxycodone (**Figure S3 B**) than middle aged males (N=7). The ANOVA confirmed significant effects for oxycodone [Sex: F (1, 10) = 13.79; P<0.005, Dose: F (1.512, 15.12) = 17.76; P<0.0005, Interaction: F (1.512, 15.12) = 8.73; P<0.01]. The Tukey post-hoc tests limited to orthogonal comparisons confirmed significant sex differences for specific doses (0, 30, 60 μg/kg/infusion) and differences within the female group for saline versus 150-300 μg/kg/infusion, as well as between either 30 or 60 μg/kg/infusion and each of 150 and 300 μg/kg/infusion.

<u>*Age differences by sex:*</u>

The middle aged <u>male rats</u> acquired fewer infusions of oxycodone than did the early adult initiating group (**Figure S3 C**) in the dose substitution experiment, as confirmed by a significant effect of Age of initiation [F (1, 13) = 10.42; P=0.01), Dose [F (1.443, 18.40) = 9.02; P<0.005] but not the Interaction. The Tukey post-hoc test limited to orthogonal comparisons confirmed a significant age-associated difference at the 30 μg/kg/infusion dose. There were also significant differences confirmed between 30 μg/kg/infusion and 150 μg/kg/infusion as well as between 60 μg/kg/infusion and each of 30 and 150 μg/kg/infusion in the early adult initiating group.

The <u>female rat</u> groups also exhibited significant differences in oxycodone IVSA as confirmed with significant effect of Age of initiation [F (1, 9) = 16.62; P<0.005), Dose [F (2.262, 20.36) = 22.61; P<0.0001] but not the Interaction. (**Figure S3 D**). The Tukey post-hoc test limited to orthogonal comparisons confirmed a significant age-associated difference at the 30-300 μg/kg/infusion doses. There were also significant differences confirmed between 60 μg/kg/infusion and 300 μg/kg/infusion conditions, as well as between 30 μg/kg/infusion and each of 60 and 300 μg/kg/infusion in the early adult initiating group. Significant differences in infusions earned were also confirmed between saline and 150-300 μg/kg/infusion as well as between either 30 or 60 μg/kg/infusion and each of 150 and 300 μg/kg/infusion in the middle aged female rats.

**Fentanyl Dose Substitution**

*<u>Sex differences in early adulthood:</u>*

The self-administration of fentanyl did not differ between male (N=8) and female rats (N=6) in early adulthood (**Figure S4 A**) and there was only an effect of dose confirmed [F (1.193, 13.72) = 14.42; P<0.005]. The post-hoc test confirmed significant differences in infusions obtained between 0.625 and 1.25 μg/kg/infusion and each of vehicle, the 2.5 and 5.0 μg/kg/infusion conditions; the latter two also differed significantly.

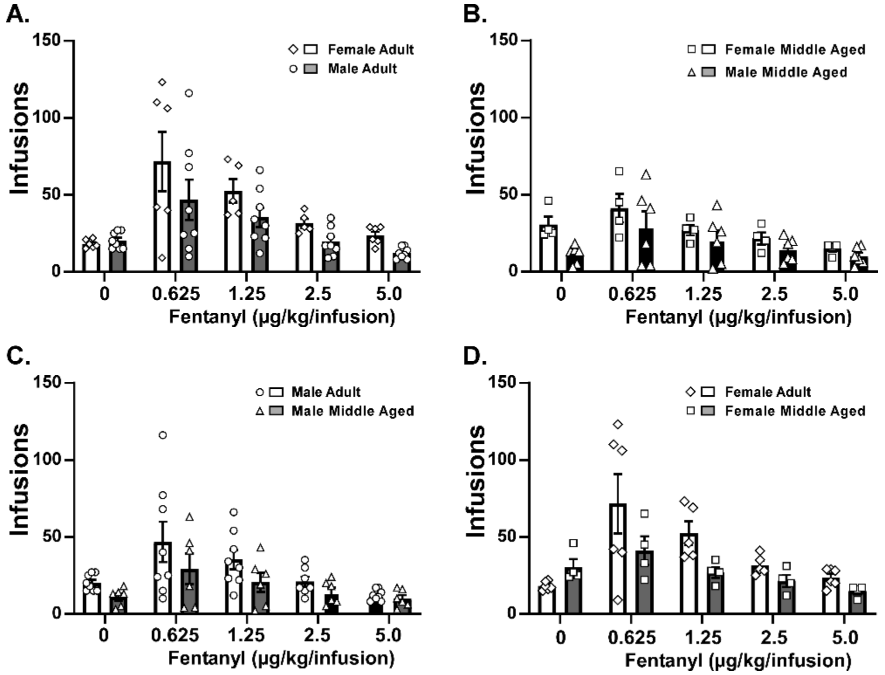


***Figure S4:*** *Mean (±SEM) and individual infusions of fentanyl (0-5 μg/kg/infusion) obtained in a dose-substitution procedure under a FR1 contingency by* ***A)*** *female and male rats which initiated IVSA at 29-30 weeks of age,* ***B)*** *female and male rats which initiated IVSA at 53-54 weeks of age, and a contrast of the male and female* ***C)*** *adult and* ***D)*** *middle aged initiating groups.*

*<u>Sex differences in middle age:</u>*

Middle aged female (N=4) and male (N=6) rats did not differ in the self-administration of fentanyl (**Figure S4 B**), but there was a significant effect of dose confirmed by the ANOVA [F (1.649, 13.19) = 8.36; P<0.01]. The post-hoc test of the marginal means confirmed significant differences between 5.0 μg/kg/infusion and each of the 0.625, 1.25 and 2.5 μg/kg/infusion conditions, as well as between 0.625 and 2.5 μg/kg/infusion.

*<u>Age differences by sex:</u>*

There were no effects of Age of initiation in the fentanyl dose response confirmed for the <u>male rats</u>

(**Figure S4 C**), however there was a significant effect of Dose [F (1.325, 16.89) = 10.20; P<0.005]. The Tukey post-hoc comparing the marginal means confirmed that significantly more infusions of the 0.625 dose were obtained compared with vehicle, 2.5 or 5.0 μg/kg/infusion, more of 1.25 μg/kg/infusion compared with vehicle, 2.5 or 5.0 μg/kg/infusion and more at 2.5 μg/kg/infusion compared with 5.0 μg/kg/infusion.

There were no effects of Age of initiation in the fentanyl dose response confirmed for the <u>female rats</u> (**Figure S4 D**), however there was a significant effect of Dose [F (1.215, 9.115) = 7.57; P<0.05]. The Tukey post-hoc test of the marginal means confirmed significant differences between 5.0 μg/kg/infusion and each of the 0.625, 1.25 and 2.5 μg/kg/infusion conditions.

#### Progressive Ratio

Only five female rats of the early adult initiating group completed the heroin dose series and four completed the oxycodone and fentanyl series; of these only two contributed to every fentanyl dose condition. Seven males of the early adult initiating group completed all three does series under the PR contingency. No effects of either sex or dose were confirmed for breakpoint in the three dose substitutions under PR for the early adult initiating groups. Nor were any significant effects on infusions obtained confirmed for heroin, oxycodone or fentanyl.

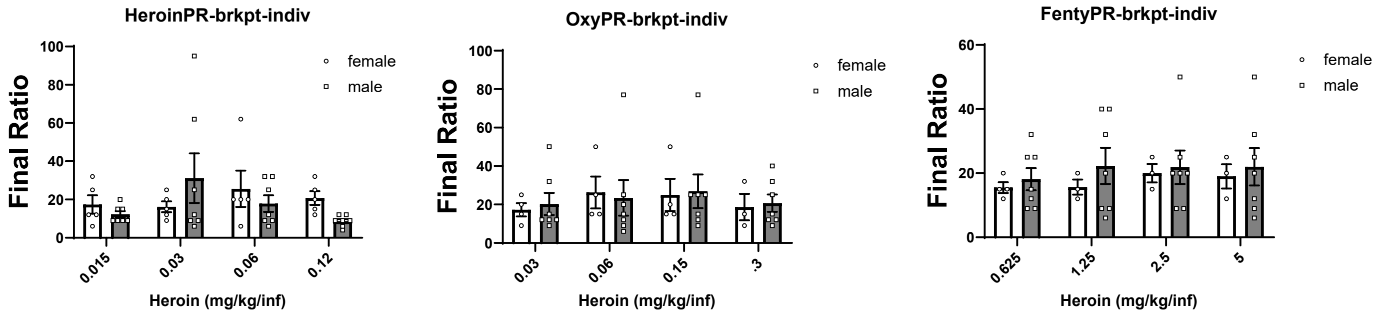


#### Supplemental Discussion

One methodological issue for the field at large to consider is the per-infusion dose when contrasting animals that differ in weight because of sex (mean female weight was ~ 48-51% of mean male weight throughout adulthood) or age, e.g, the mean male bodyweight was ~20% higher, and the mean female bodyweight was ~7-10% higher, in 1yr olds compared with the weight of the earlier initiating groups during IVSA. Sex-related differences in *brain size* of rats are, however smaller since female rat brains weigh 90-95% of male brain weight. Drugs distribute differentially to body tissues on different time-courses thus selection of an ideal i.v. dose to produce equivalent brain exposure is complicated (Volkow et al., 2010). In this study, mean heroin infusions in the earlier initiating males were 47.5% of the female group mean at the end of acquisition. Mean heroin infusions were 45-62% of the female amount across the dose substitutions under the FR1 schedule. In contrast, mean heroin infusions obtained by the middle aged males was about 48% of the younger male group mean at the end of acquisition. Likewise, the older initiating group of males' mean infusions earned in the FR1 for heroin was 42%-51% of the younger initiating animals' means. This suggests the age-associated difference is not entirely explained by the 20% higher bodyweight, although brains grow in adult rats and reach peak weight and volume around middle age (Zhao and Zahr, 2025). This may be a general unresolved issue in the field, since most IVSA procedures similarly adjust doses across time and sex by bodyweight and not by brain weight.

Popular media descriptions of fentanyl-related harms often emphasize the potency of this drug relative to oxycodone and heroin. This is often stated without reference to the assay from which the estimate is derived and in any case potency relationships may not be the same for, e.g., analgesic versus reinforcing properties. In this study, the dose-substitution in the same group of animals estimated that fentanyl is approximately 6 times more potent than heroin. That is, early adult initiating female rats obtained 23.7 infusions of 5 μg/kg/inf fentanyl (118.5 μg/kg session total) and 25.0 infusions of 30 μg/kg/inf heroin (750 μg/kg session total), suggesting a rough behavioral equivalency at these per-infusion doses and therefore a 6.3-fold potency relationship. Similarly, the early adult initiating male rats obtained a mean of 21.5 infusions of the 15 μg/kg dose of heroin (322.5 μg/kg session total) and a mean of 21 of the 2.5 ug/kg dose of fentanyl (52.5 μg/kg session total) and thus again, a 6.1-fold potency relationship for overall session intake.
